# Exploring the Inner Workings of Neuron Circuits That Exhibit Persistent Activity To Explain How Working Memory and Executive Function Are Implemented in The Brain

**DOI:** 10.1101/2021.07.05.451167

**Authors:** Paul Gomez, Jeremy Gomez, Andres Gomez

**Author notes:** **Correspondence:** Dr. Paul Gomez, Nanobiotek LLC, Pembroke Pines, Florida, USA.

## Abstract

In this research we explore in detail how a phenomenon called “sustained persistent activity” is achieved by circuits of interconnected neurons. Persistent activity is a phenomenon that has been extensively studied (Papoutsi et al. 2013; Kaminski et. al. 2017; McCormick et al. 2003; Rahman, and Berger, 2011). Persistent activity consists of neuron circuits whose spiking activity remains even after the initial stimuli are removed. Persistent activity has been found in the prefrontal cortex (PFC) and has been correlated to working memory and decision making (Clayton E. Curtis and Daeyeol Lee, 2010).

We go beyond the explanation of how persistent activity happens and show how arrangements of those basic circuits encode and store data and are used to perform more elaborated tasks and computations.

The purpose of the model we propose here is to describe the minimum number of neurons and their interconnections required to explain persistent activity and how this phenomenon is actually a fast storage mechanism required for implementing working memory, task processing and decision making.

## INTRODUCTION

In this research we explore the existence of neuron circuits in the brain that store information in a way that has not been well understood. It is known that memories are kept in the brain in the synapses of large collections of neurons (Mayford et al. 2012). These synapses are dynamically modified through learning processes and adaptation. Synaptic plasticity is accomplished by two opposite processes: long-term potentiation (LTP) and long-term depression (LTP). Changing the strength of a synapse by means of LTP or LDP takes between 10 and 30 minutes in average (Wang X. et al. 2015; Rosenberg et al. 2016; Le Duigou et al. 2015; De Pasquale et al. 2014; Wang Hui et al. 2016).

To perform intermediate calculations in a timely manner, we hypothesize that the brain also keeps memories on neuron arrangements like the one proposed in this research article. The information stored on such circuits store information as sustained bursts of action potentials that can be activated (set) or deactivated (reset) on demand, thus, storing one bit of information at speeds several orders of magnitude faster than the long-term memories stored on synapses. The brain needs to react quickly to external stimuli and to do so we argue that neuron flip-flops are used to provide this type of immediate temporary memory. We introduce the term Neuron Flip-Flop which is a circuit made of three neurons connected in such way that persistent activity is achieved. We name these circuits neuron flip-flops (NFF) for their similarity to equivalent devices in the computer engineering world (Gupta and Mehra, 2016). Neuron flip-flops also facilitate sequential processing which is a way to execute algorithms in the brain. Neuron flip-flops hold intermediate states during a multi-step task. Those states can be interrogated and decision making is made by evaluating the current state of a neuron flip-flop (NFF).

These neuron circuits (NFF) have been found in the prefrontal cortex (PFC) (Compte Albert, 2006; Fellous et al. 2003) and could also be probably found in the CA1 or CA3 nuclei of the hippocampus where endogenous oscillations have been found (Jochems et al. 2013; Pinsky et al. 1994; Xu et al. 2008), the hippocampus entorhinal cortex (EC) and amygdala (Egorov et al., 2006) and very likely on the dentate gyrus (DG) which has neurons whose spiking pattern is of type NASP (Non-Adapting Spiking) as classified by the Hippocampome project (Komendantov et al. 2019; Hippocampome). These DG neurons exhibit the closest spiking pattern matching the results of our simulations.

It is worth mentioning that oscillations (bursts of spikes) are present in the brain and have been studied extensively. As an example, pyramidal CA3 neurons in the hippocampus exhibit endogenous bursts of spikes whose frequency has been found to be associated with the intensity of the stimuli (Balind et al. 2019; Harris et al. 2001). Theta oscillations are widely found in the hippocampus and associated to several functions. For example, neocortical neurons are significantly phase-locked to hippocampal theta oscillations during either spatial exploration or REM sleep (Sirota et. al. 2008). Theta oscillations are most regular in frequency and largest amplitude in the stratum lacunosum-moleculare of the hippocampal CA1 region and are also present in the dentate gyrus and the CA3 region (Buzsa 2002). Theta oscillations have also been found in the amygdala (Paré D et al. 2000) and in the entorhinal cortex (Alonso A. et al. 1987).

Based on our hypothesis that a different memory storage mechanism must exist in the brain, we devised a model that has three neurons interconnected in such way that can store one bit of information without resorting to the plasticity of their synapses. What this circuit stores is an oscillation (a burst of action potentials) and the oscillate or do-not-oscillate state is controlled on demand.

The neuron circuit proposed in this research is made of three neurons that do not necessarily exhibit endogenous bursting features as those found in CA3 pyramidal neurons.

The circuit, named Set-Reset Neuron Flip-Flop (SR NFF), has two independent inputs (a Set signal and a Reset signal) and one output, the Q signal that represents the value stored on the circuit.

Figure 1 shows the SR NFF circuit along with the timing of the input, feedback and output signals. The Set signal is a presynaptic action potential that is applied to a dendrite having an excitatory synapse on input neuron X0. The arrival of an impulse on this input dendrite will drive all three neurons in a sustained spiking state that remains oscillating even after the stimulus is removed, thus storing a value on the circuit (bursting).

**Figure 1.**
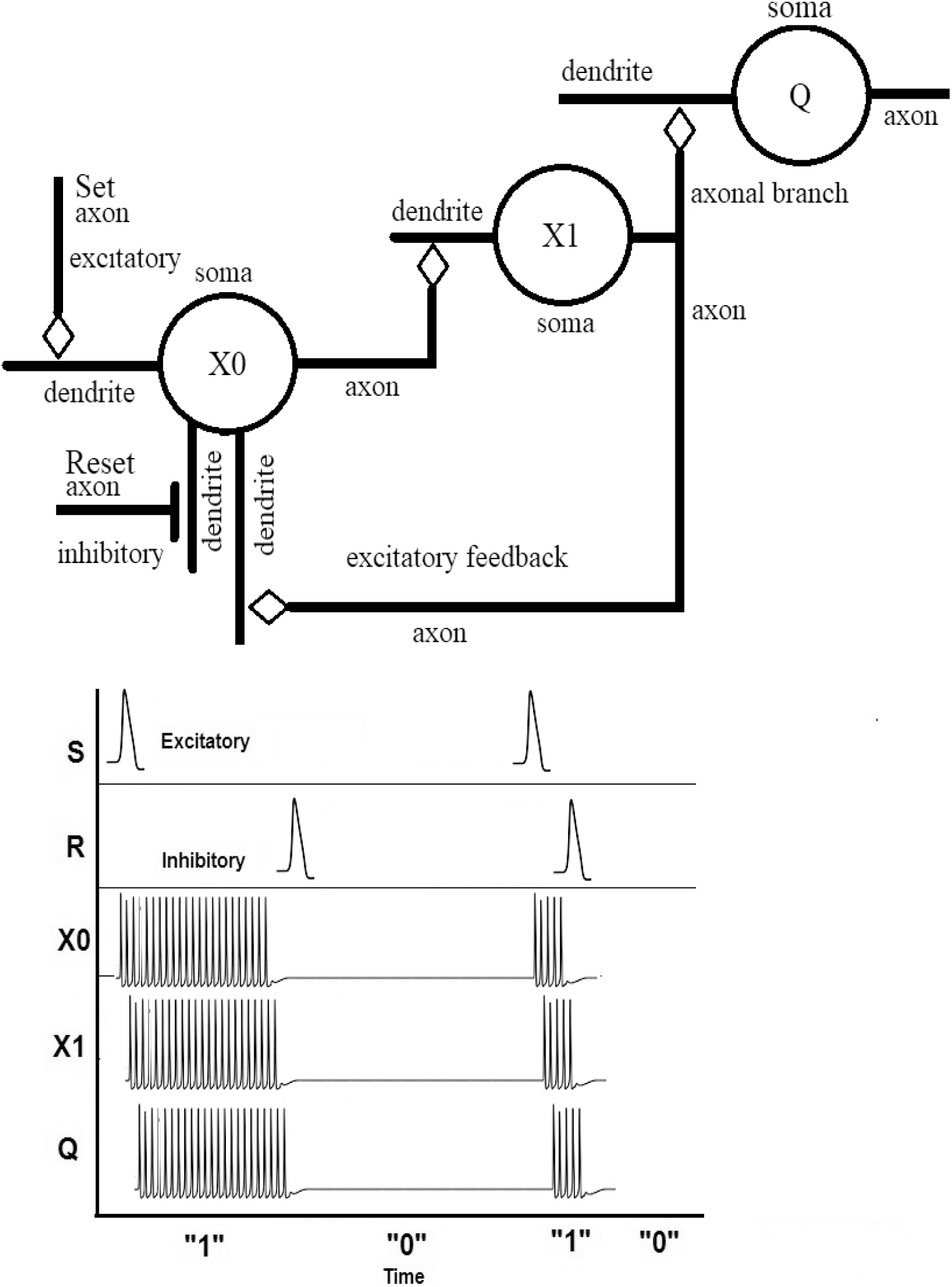
A Neuron Set-Reset Flip-Flop

Output neuron (Q) is not required for the persistent activity (sustained spiking) but it is necessary to deliver the value of the device to other neuron circuits. Neuron Q must have dozens or even hundreds of axon terminals and must be capable of sending neurotransmitters to them at a fast pace. Neurons X0 and X1 on the other hand, only need an axon with a single axon terminal and one axonal branch.

The oscillation is carried out by the positive feedback connection between middle neuron X1 axon and the feedback dendrite of input neuron X0. This feedback dendrite has an excitatory synapse with axon X1 terminal. Figure 4 shows the sequence of events; once the input stimulus makes neuron X0 fire, an action potential travels on axon X0 to input dendrite of middle neuron X1, which arrives at the soma of neuron X1 which in turn fires an action potential on its axon. As the action potential on axon X0 dies, an action potential from X1 arrives at its feedback dendrite resulting in neuron X0 fire a new action potential. The cycle is repeated indefinitely until an action potential arrives at Reset dendrite of input neuron X0. The reset dendrite has an inhibitory synapse that causes hyperporalization of the soma of X0.

The spiking / no spiking state of the circuit effectively stores one bit of information. It is important to emphasize that the input stimuli can be removed and the circuit will stay in the state determined by the nature of the stimulus (Set/Reset).

The output neuron Q relays the sustained bursting to its axon which is the output of the circuit. One axonal branch of axon X1 with an excitatory synapse on a dendrite of neuron Q transmits the signal from X1 to Q.

The period of the oscillation is the sum of time delays due to the geometry of dendrites and axons (length and diameter), impedances, capacitances, and synapses delays. There are not specific required parameters to have this circuit behave in this way. Several other configurations are possible, such as all signals -Set, Reset and Feedback- connected to a single dendrite on neuron X0, or even axo-somatic or axo-axonic synapses as well.

To test the circuit’s physiological ability to function as proposed, a model was developed using software tool Neuron© by Yale University. The model and its results are explained in the next section.

## SIMULATION OF A SINGLE NEURON FLIP-FLOP

Software tool Neuron© by Yale University (Carnevale, N.T. and Hines, M.L 2006) was used to create a model according to Figure 1. The parameters for all somata, dendrites and axons were defined with the default values Neuron© provides. Only dendrite and axon lengths were specified and synapses were tuned to exhibit the required inhibitory or excitatory feature as well as time delays.

The resulting data was exported to external files and MATLAB® was used to read the files and generate the figures presented below.

To simulate the Set and Reset signals, current clamps were used at different times. The currents are injected on input neuron X0 respective dendrites. The length of the experiment was 500 ms. Figure 2 shows an excitatory current applied to neuron X0 Set dendrite at times 10 ms and 350 ms, and an inhibitory current applied at times 250 ms and 400 ms on Reset dendrite. The spiking voltage of axon X0 is shown immediately after the Set signal is applied. Axon X0 stops spiking and stays at rest potential (polarized) as expected after a Reset signal is injected.

**Figure 2.**
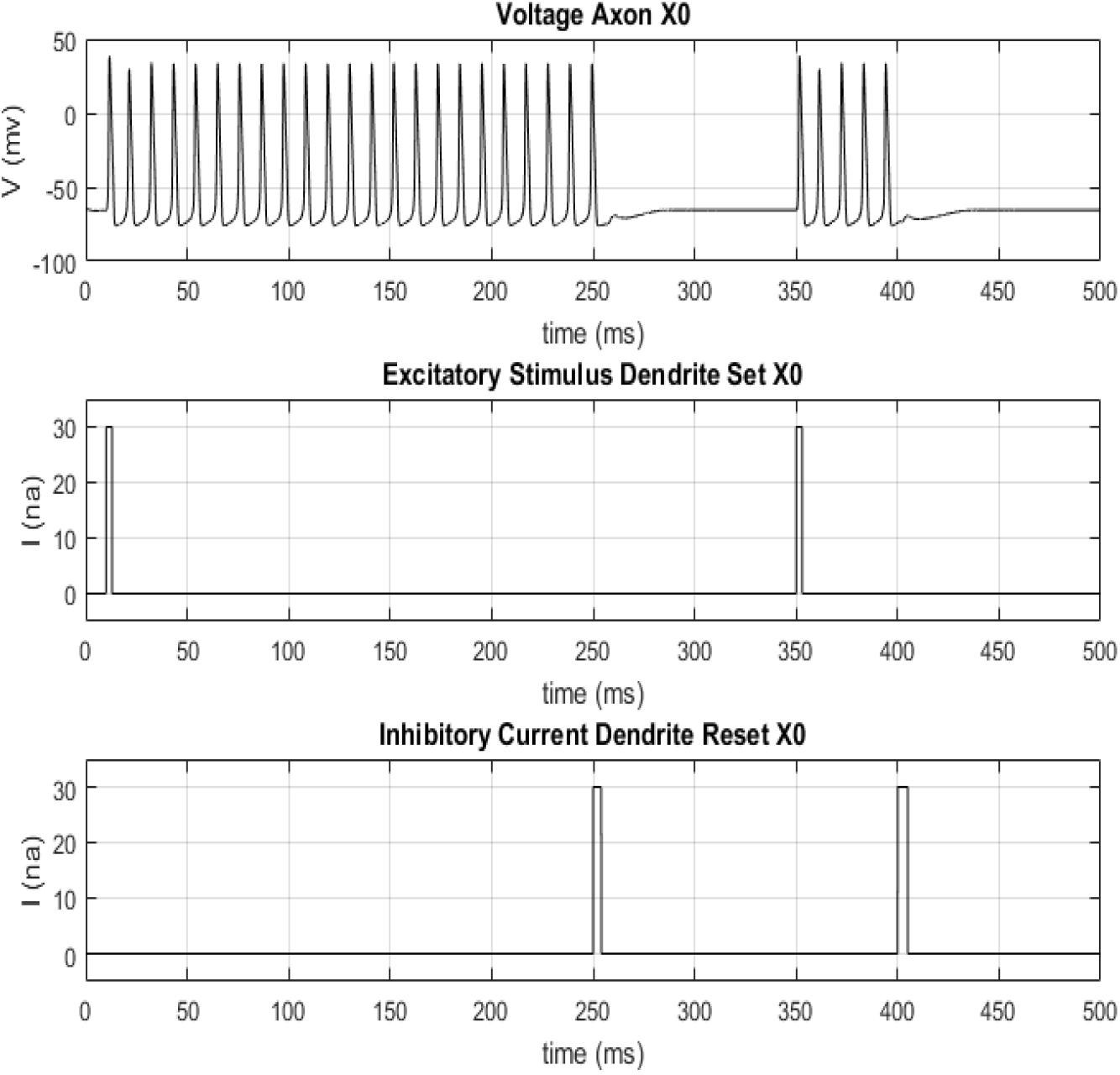
Axon X0 voltage relative to excitatory (Set) and inhibitory (Reset) signals.

Figure 3 shows the action potentials of all axons along the 500 ms. There is an inherent delay between the spikes of the three neurons due to the propagation delay down the dendrites, axons and synapses. This delay is shown in more detail on Figure 4, which shows the first 45 ms of the experiment only.

**Figure 3.**
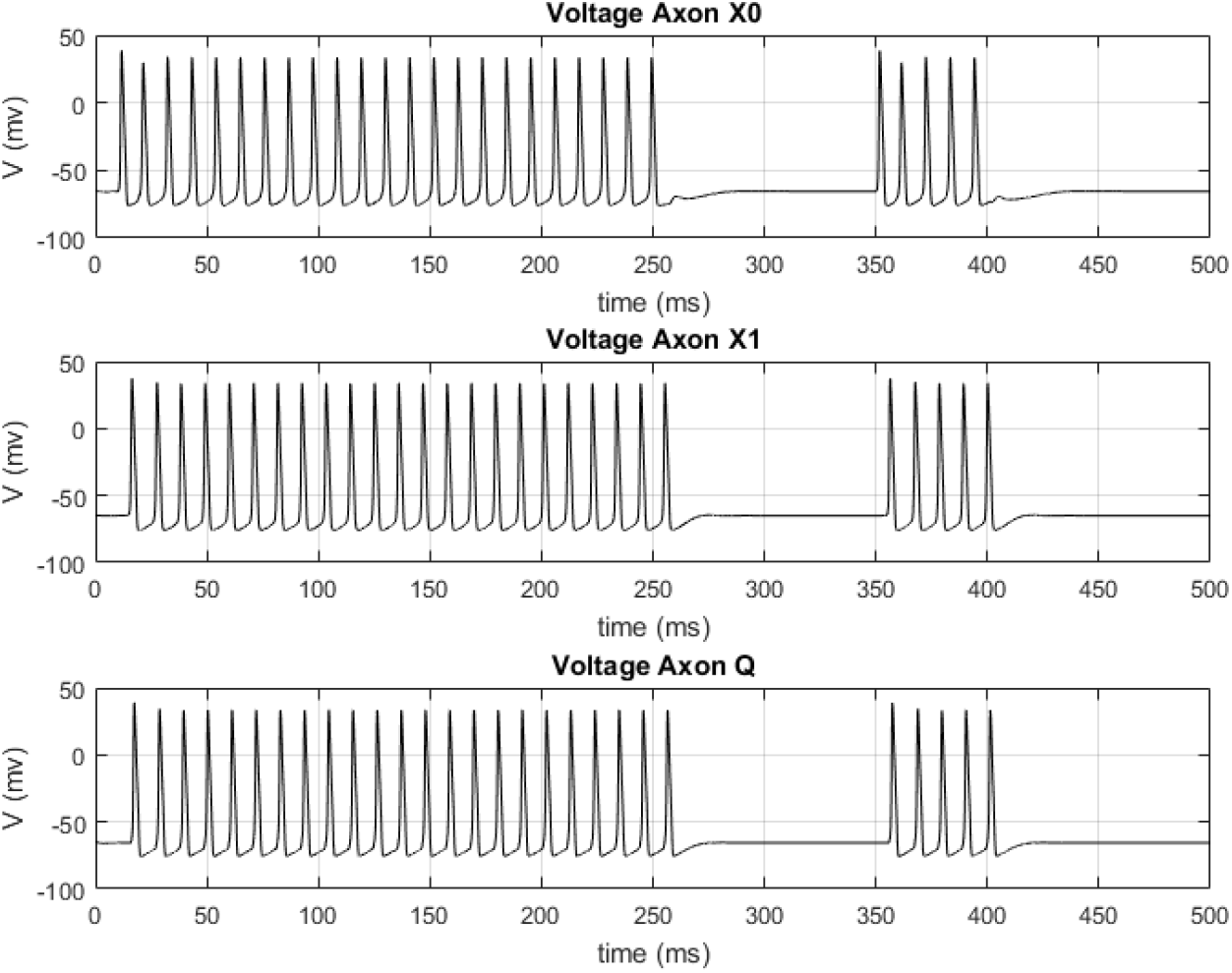
Voltages on all three axons.

**Figure 4.**
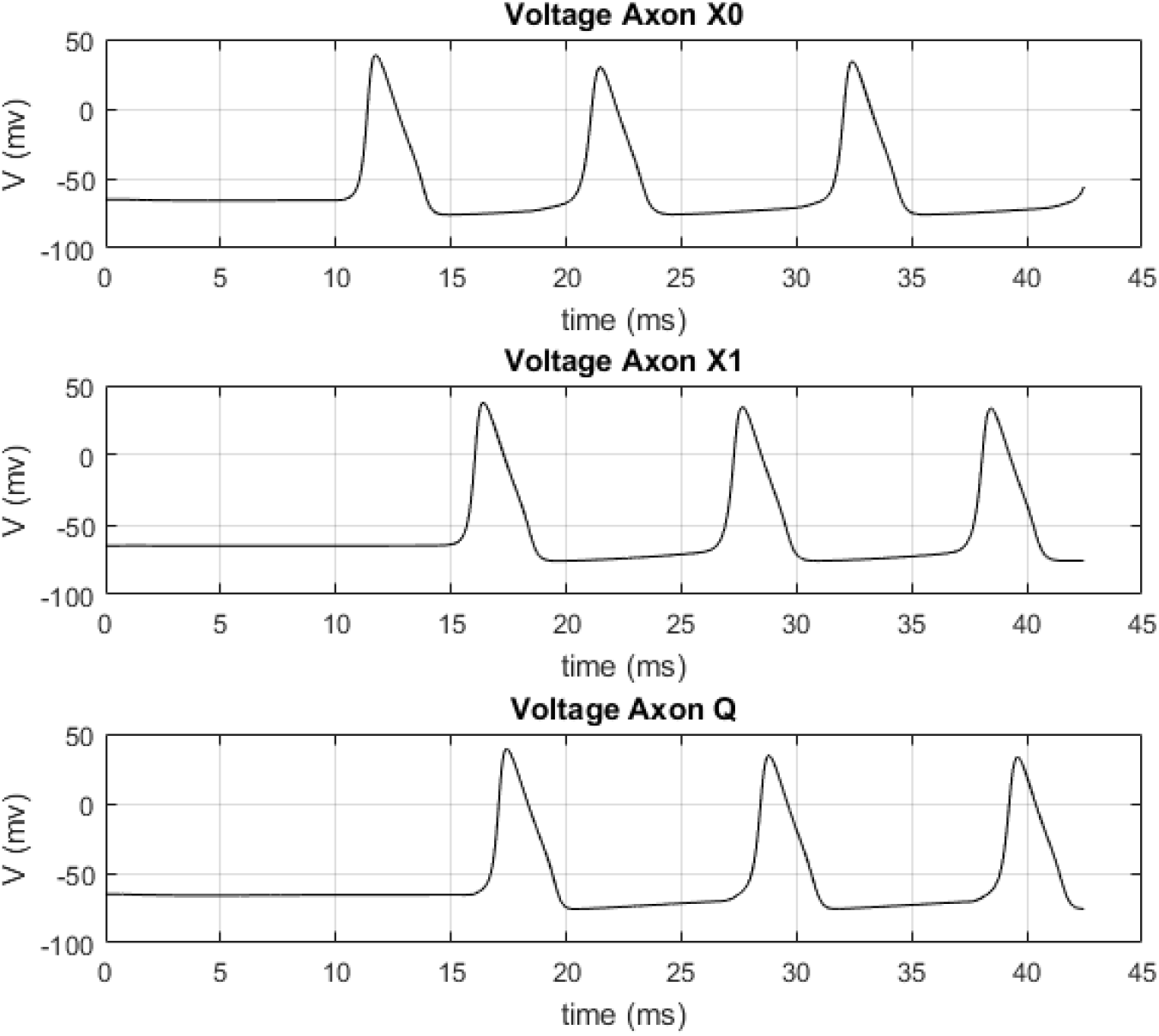
Voltages on all three axons for the first 45 ms showing the delays between them.

## NEURON FLIP-FLOPS AT WORK (WORKING MEMORY)

Working Memory refers to our ability to temporarily maintain and process information (Masse, Song et al. 2019), (Izaki et al., 2008). Having elucidated the inner workings of a circuit that exhibits sustained persistent activity and showing that effectively, this circuit stores one bit of information, we continue to show how arrays of several NFF wired appropriately implement more complex circuits that perform working memory tasks.

We have chosen a simple task we are all familiar with: perform the arithmetic sum of two one-digit numbers. Figure 5 shows schematically the components of a circuit that computes R = A + B where A and B are one digit numbers, from 0 to 9.

**Figure 5.**
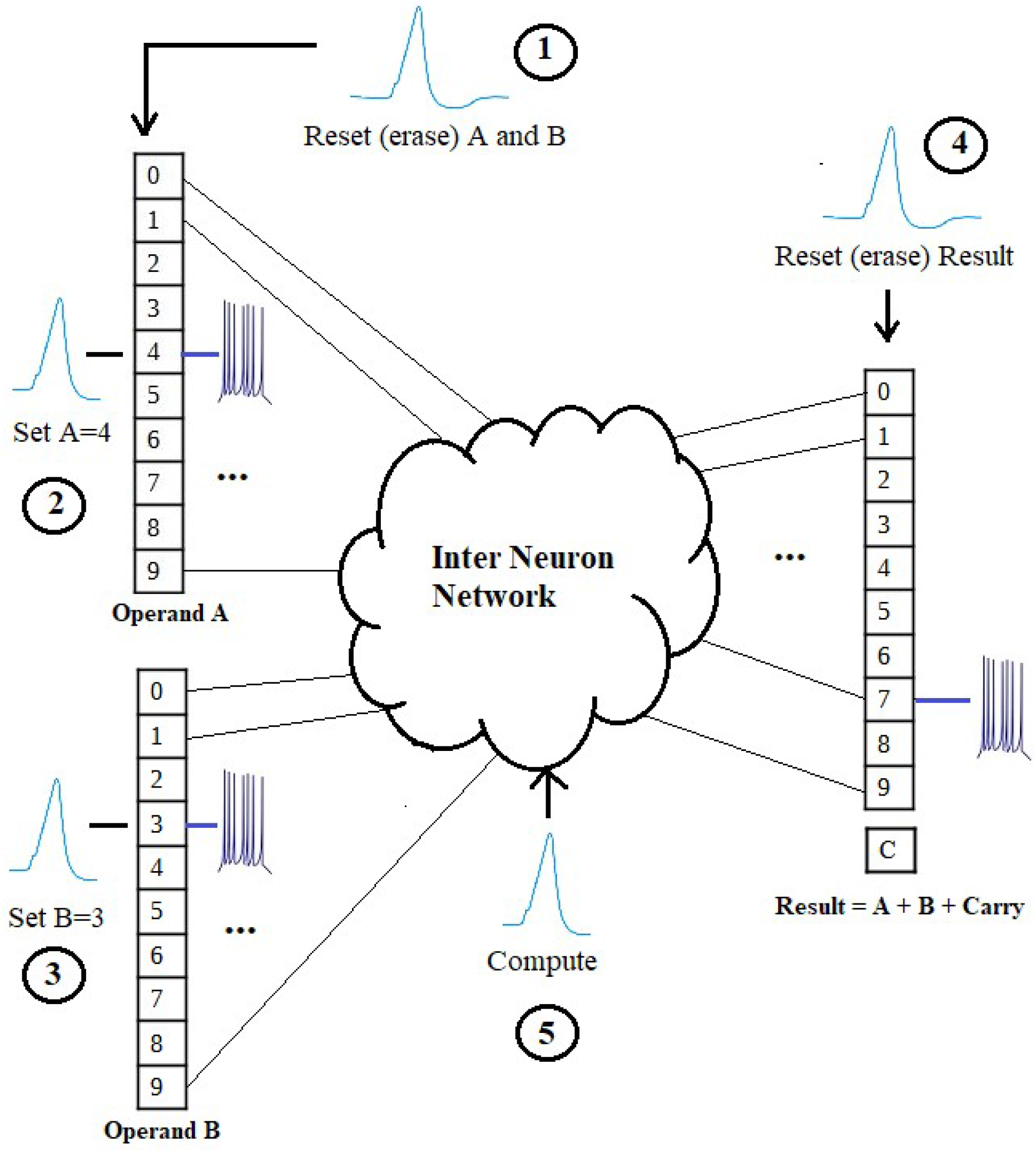
Computation of the sum A+B with 2 registers for the operands and one register for the result. The numbers in circles indicate the sequence of events.

On the left side of the figure the two operands, A and B, are shown as registers (arrays of neuron flip flops). Each number is represented as one-and-only-one flip-flop on its ON state (persistent activity) at a given time. In this example, the objective is to obtain the sum 4+3, and thus, on register (operand) A, its flip-flop # 4 is set by applying an action potential on its Set input. Similarly, for register (operand) B, its flip-flop # 3 is set by applying an action potential on its Set input. The result of the sum, 7, is obtained by means of an inter network of neurons that are wired to perform the sum (what we know as the sum fact table). This inter network has 10 outputs, one for each possible result (0-9) which delivers by appropriately sending an action potential to the appropriate Set input of the Result register.

The adder circuit was modeled and simulated with Neuron© with the values shown on Figure 5, Z = 4 + 3. On Figure 6, the values of all 10 bits of operand A are shown. Only flip-flop #4 exhibits persistent activity while others are at their resting (polarized) potential. Similarly on Figure 7, operand B value is shown, with flip-flop #3 spiking while all others are polarized. Similarly, on Figure 8, the value of the result, register Z, is shown: only flip-flop #7 exhibits sustained persistent activity.

**Figure 6.**
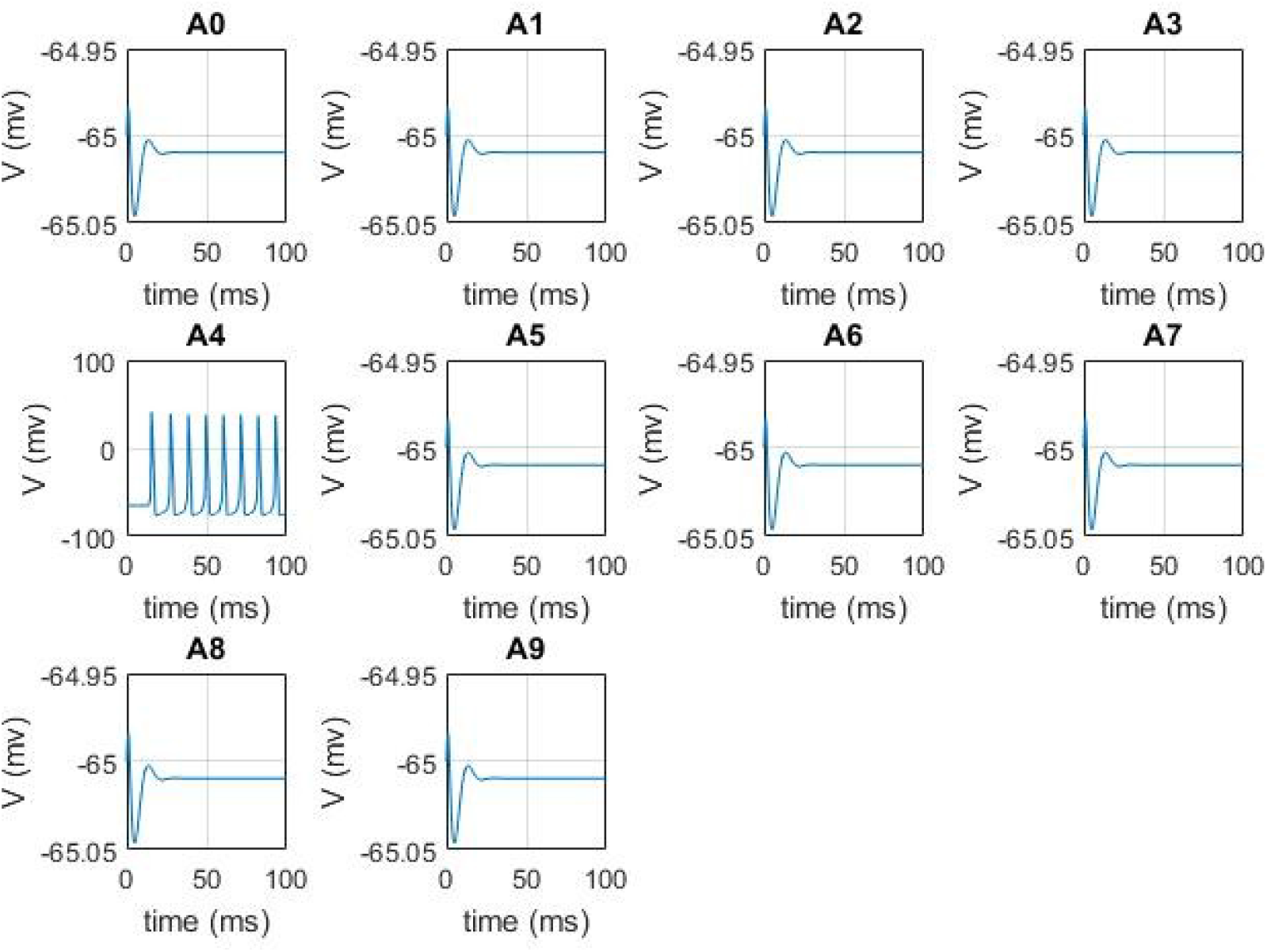
Operand A: Representation of number 4 in the brain with an array of 10 neuron flip-flops.

**Figure 7.**
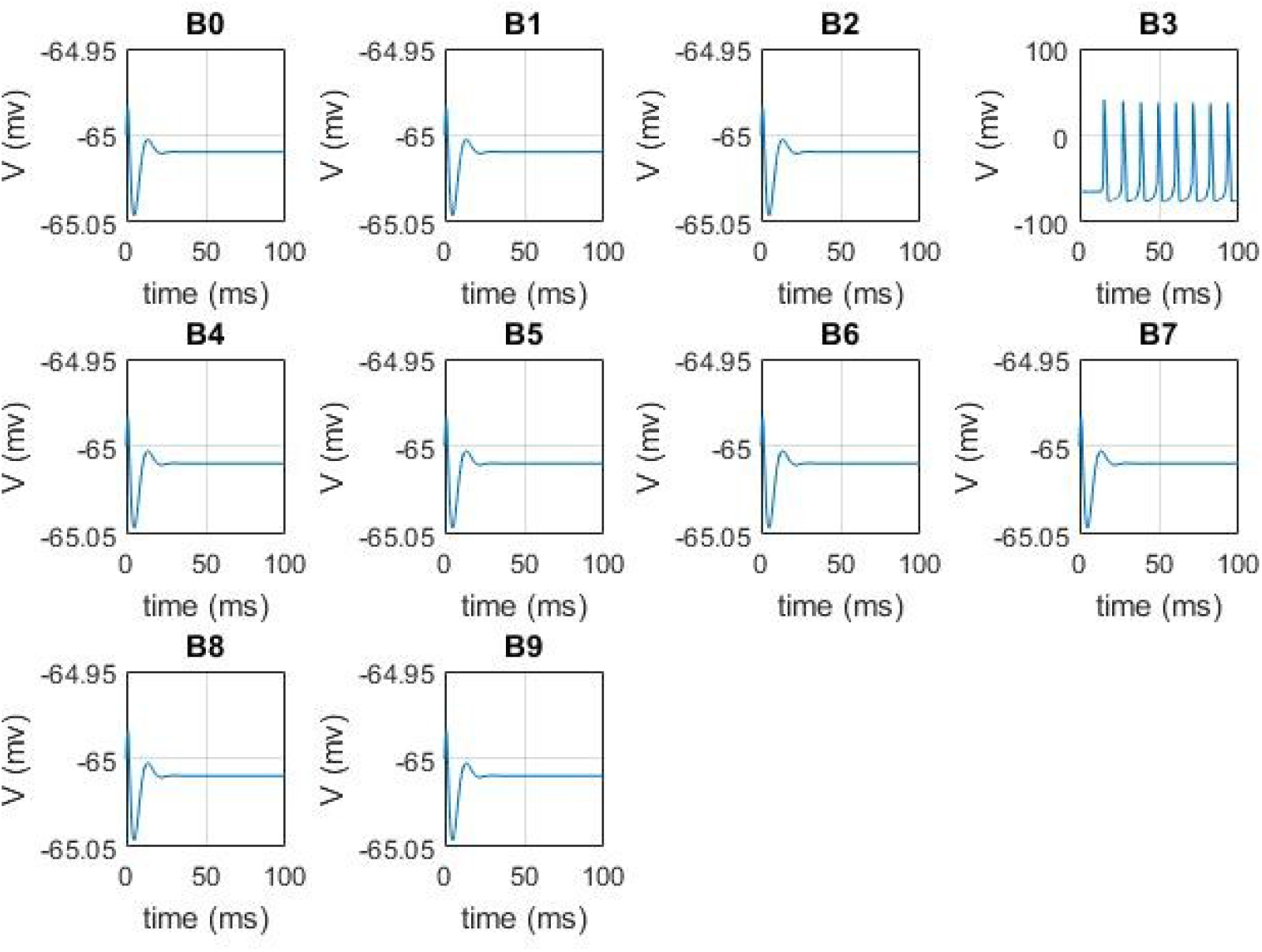
Operand B: Representation of number 3 in the brain with an array of 10 neuron flip-flops.

**Figure 8.**
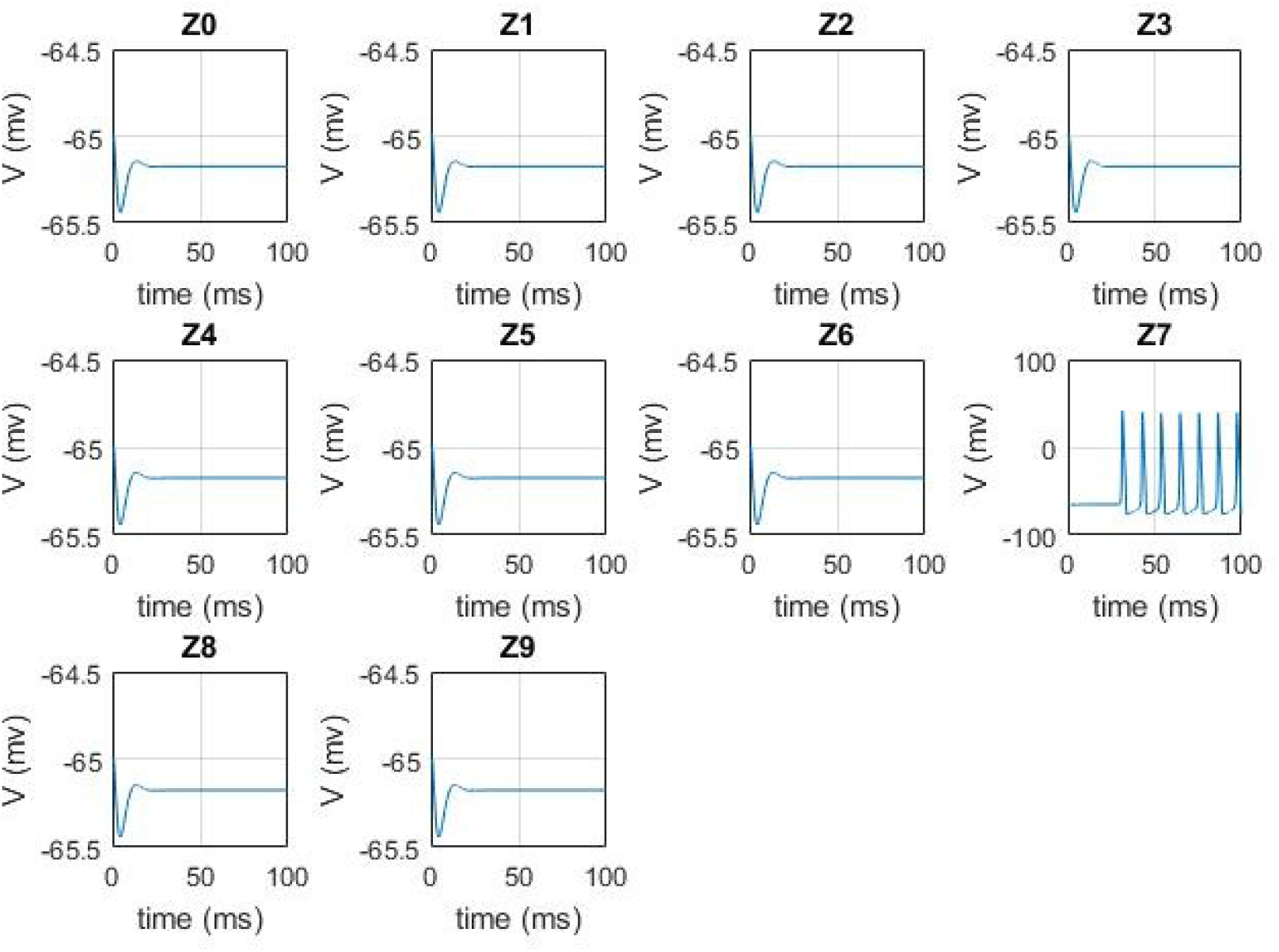
Resulting sum S: Representation of number 7, the sum of operands A and B, in the brain with an array of 10 neuron flip-flops.

Even a simple task as adding two one digit numbers, requires a series of steps, which are numbered in circles on Figure 5. These are the steps:

1. Reset (erase) all neuron flip-flops of registers (operands) A and B by applying an action potential to all the Reset inputs of all flip-flops
2. Load value of operand A: Apply an action potential to the Set input of the appropriate flip-flop of Register A.
3. Load value of operand B: Apply an action potential to the Set input of the appropriate flip-flop of Register B.
4. Reset (erase) all neuron flip-flops of register Result
5. Compute. This is an action potential that sets a flip-flop that is connected in a logical AND way which each of the 10 output lines of the Inter Neuron Network. Only one is transferred to the appropriate Set input of register Result, thus, yielding the sum of A and B.

Now, the next obvious question and a very important one is, how the brain implements sequence?

The answer is again, a variation of the neuron flip-flop (NFF) called a monostable (as they are known in the computer engineering world) (Chien Hung-Chun and Lo Yu-Kang, 2011). These are circuits that hold one bit of information for some predefined time and reset themselves after the time is up. The following section explains in more detailed the circuits that implement task sequencing in the brain.

## EXECUTIVE FUNCTION: TASK SEQUENCING

We argue that neuron flip-flops are the fundamental circuits (devices) in the brain that implement fast temporary storage (working memory) and also, with some additional interconnections, provide the function of task sequencing, necessary to perform conscious tasks that require several steps executed in sequence.

On Figure 9 we show the basic monostable circuit, which consists of a neuron flip-flop with a delay line connected from its output, Q, to its own reset input. A delay line is simply implemented by several neurons connected in daisy chain. The time required for an action potential to arrive at the reset input, is the delay of the monostable. This circuit, once an action potential is applied to its Set input, begins spiking on its output axon Q, exhibiting persistent activity. However, the output axon Q, has a synapse with a neuron which in turn has its axon connected to another neuron’s dendrite, perhaps several neurons in cascade, in such way that they implement a delay line that has its last axon connected to the Reset input of the neuron flip-flop. That way, after some time determined by the number of neurons on the delay line and their physical characteristics, the neuron flip-flop resets itself.

**Figure 9.**
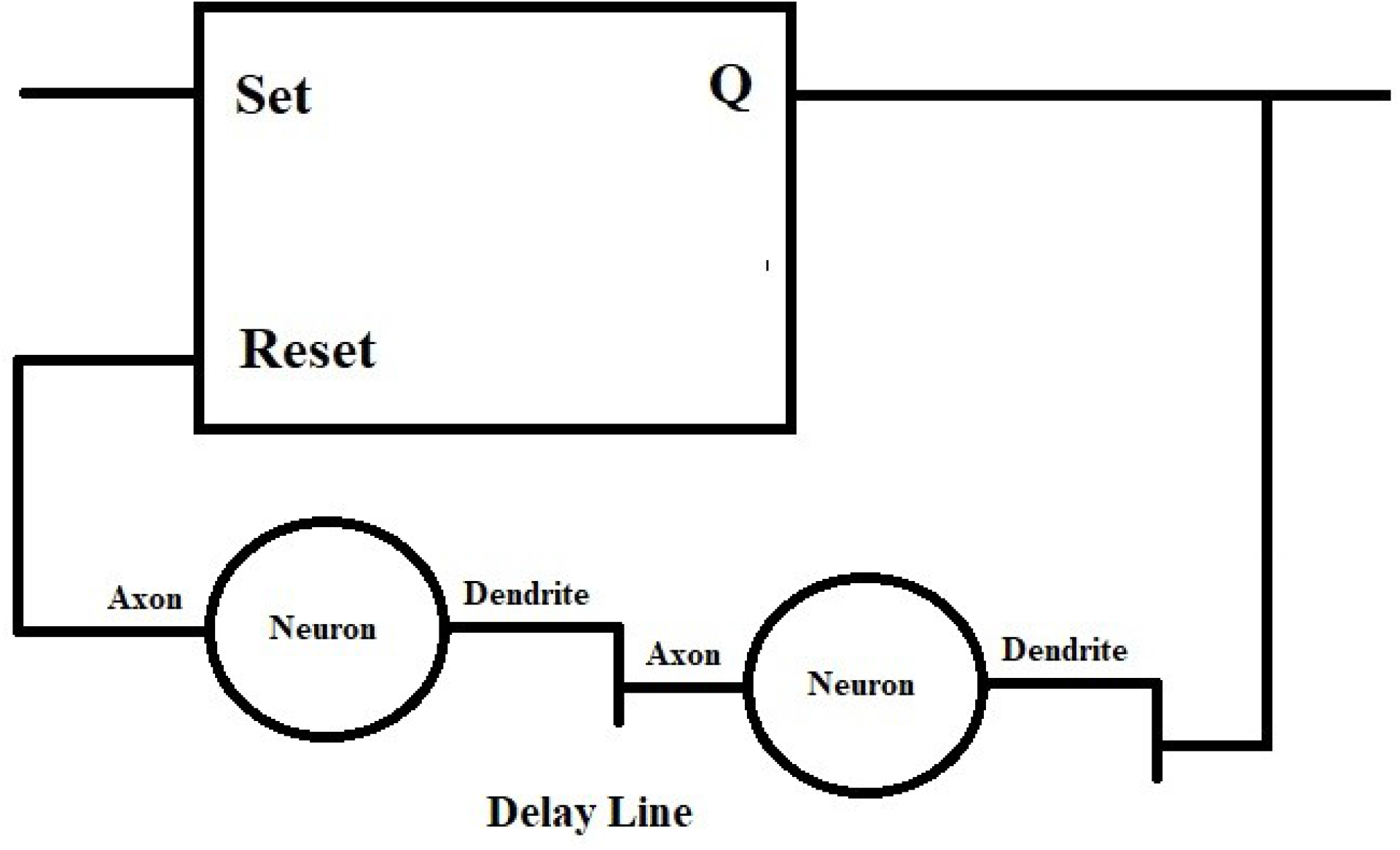
Monostable Circuit. The neuron flip-flop resets itself after a delay time. The delay line consists of several neurons connected in daisy chain.

By connecting several monostables in sequence as shown on Figure 10, a task sequencer is effectively implemented. The output axon action potential of each stage is used to activate another neural circuit that performs a particular task, thus, a task sequencer circuit executes several steps sequentially.

**Figure 10.**
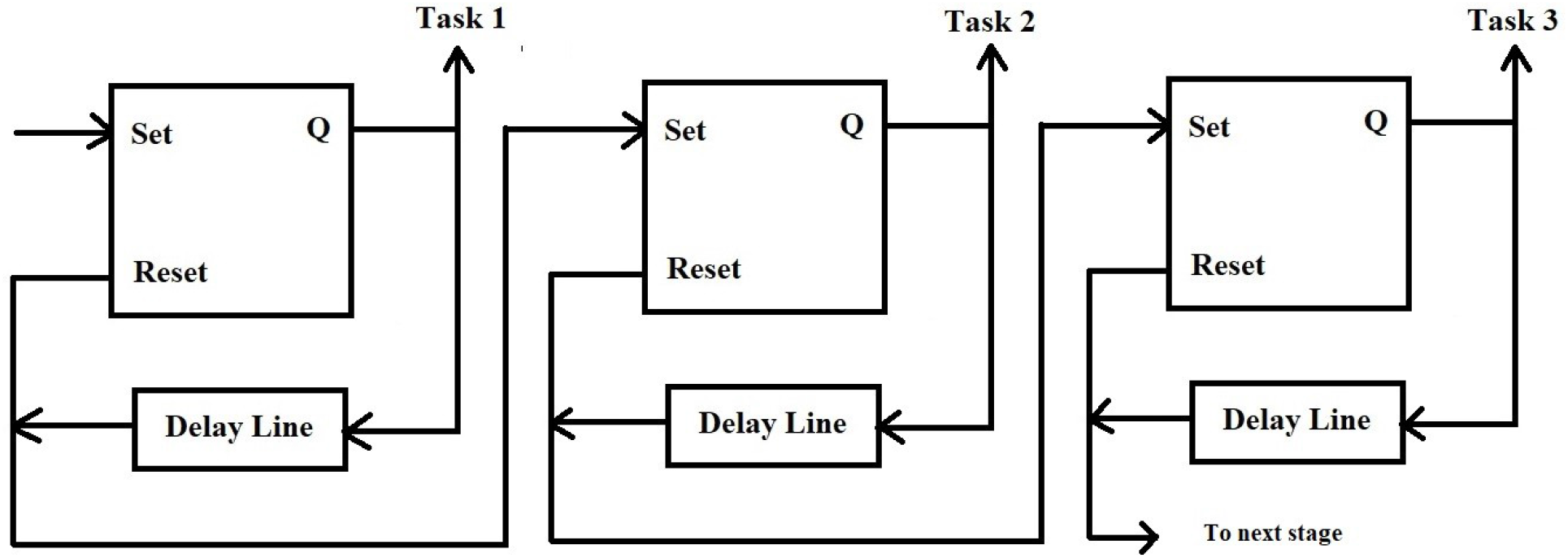
Task Sequencer Circuit

The circuit was simulated with Neuron© and the resulting signals are shown on Figure 11.

**Figure 11.**
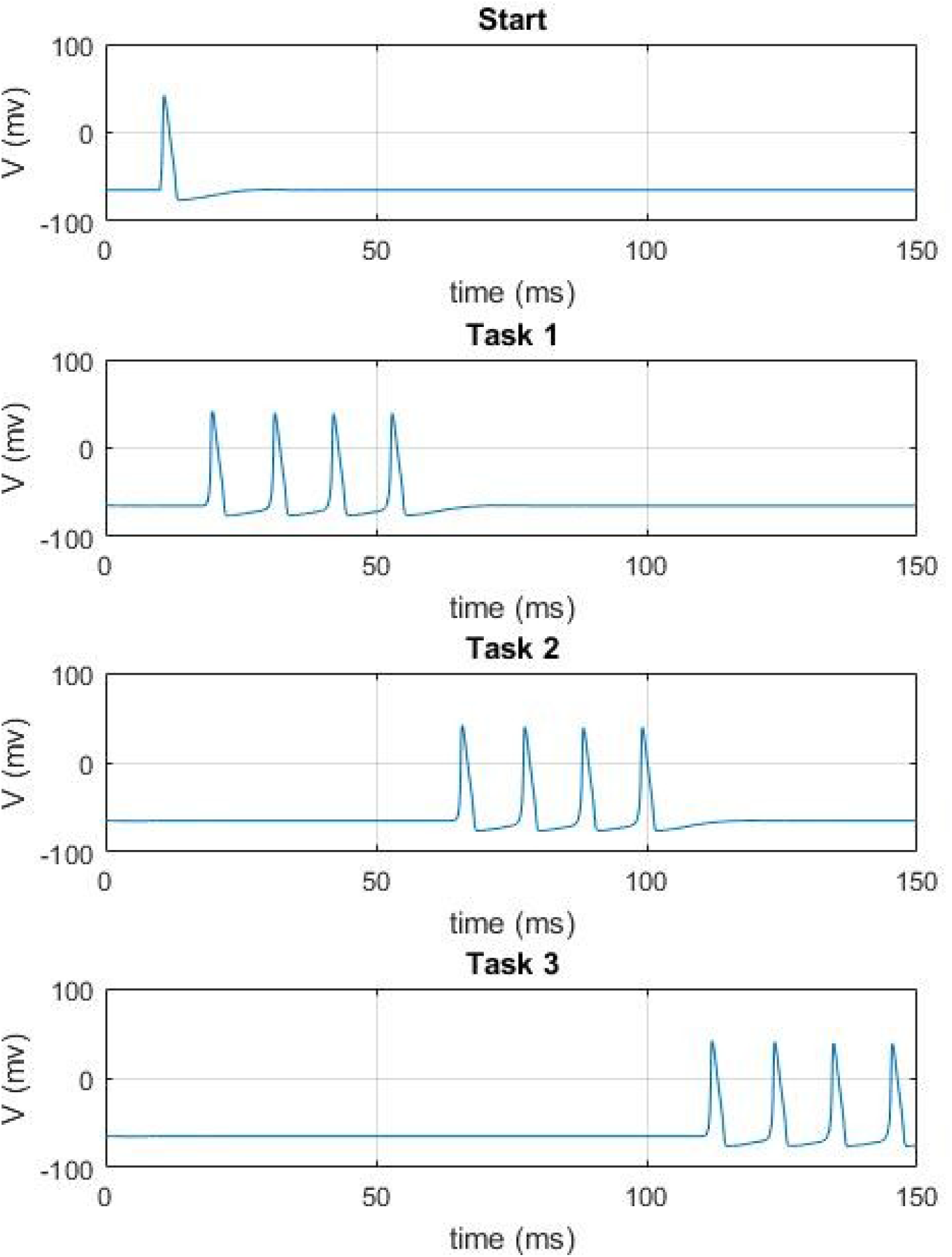
Sequencer Simulation.

## CONCLUSION AND FUTURE WORK

The brain needs to perform computations that require fast, intermediate storage that cannot be implemented on slow adapting synapses. Additionally, the brain executes complex executive function tasks in sequences of several steps. We hypothesize that all these functions are based on a well known phenomenon called Sustained Persistent Activity. A neuron flip-flop circuit that stores one bit of information has been proposed, modeled, simulated and the results are found to align with the expectations. Arrays of neuron flip-flops are used by the brain to symbolically represent data and arrangements of neuron flip-flops are used to implement task sequencing.

Future research will continue to determine how other critical functions take place in the brain, namely, how comparing two values is accomplished; how the result of a comparison is used to branch to another step in a task sequence; how data is moved between arrays of neuron flip-flops and between arrays of NFFs and other neural networks on the neocortex and other areas, which hold memories on their synapses.

## AUTHOR CONTRIBUTIONS

PG conceptualized the idea for the research and supervised the project. All authors developed the concepts, performed Neuron© simulations and wrote the manuscript. JG wrote the code for the simulation of the monostable and task sequencer. AG and PG wrote the code to simulate the neuron flip-flop and the adder circuit All authors read and approved the final manuscript.

## ETHICAL STATEMENT

We did not perform any experiments with animals nor humans.

## FINANCIAL STATEMENT

The authors did not receive any funding for this project.

## DATA AVAILABILITY STATEMENT

The data and the Neuron© code that support the figures of this study are available from the corresponding author upon request.

## REFERENCES

Alonso A, García-Austt E. 1987, Neuronal sources of theta rhythm in the entorhinal cortex of the rat. I. Laminar distribution of theta field potentials. Exp Brain Res. 1987;67(3):493–501. doi: 10.1007/BF00247282. PMID: 3653311.

Balind Raus, S., Magó, Á., Ahmadi, M. et al. 2019, Diverse synaptic and dendritic mechanisms of complex spike burst generation in hippocampal CA3 pyramidal cells. Nature Communications 10, 1859 (2019). https://doi.org/10.1038/s41467-019-09767-w

Buzsa Gyorgy, 2002, Theta Oscillations in the Hippocampus, Neuron, Vol. 33, 325–340, January 31, 2002

Carnevale, N.T. and Hines, M.L. 2006, The NEURON Book. Cambridge, UK: Cambridge University Press, 2006.

Chien Hung-Chun, Lo Yu-Kang, 2011, Design and implementation of monostable multivibrators employing differential voltage current conveyors, Microelectronics Journal Volume 42, Issue 10, October 2011, Pages 1107–1115

Clayton E. Curtis and Daeyeol Lee, Beyond working memory: the role of persistent activity in decision making, Trends Cogn Sci. 2010 May; 14(5):216–22.

Compte Albert, 2006, Computational and in Vitro Studies of Persistent Activity: Edging Towards Cellular and Synaptic Mechanisms of Working Memory, Neuroscience 135–151

De Pasquale Roberto, X Thiago F. Beckhauser, Marina Sorrentino Hernandes, and Luiz R. Giorgetti Britto, 2014, LTP and LTD in the Visual Cortex Require the Activation of NOX2, The Journal of Neuroscience, September 17, 2014 • 34(38):12778–12787

Egorov Alexei V., Unsicker Klaus and Oliver von Bohlen und Halbach, Muscarinic control of graded persistent activity in lateral, amygdala neurons, European Journal of Neuroscience, Vol. 24, pp. 3183–3194, 2006

Fellous Jean-Marc and Sejnowski Terrence J., 2003, Regulation of Persistent Activity by Background Inhibition in an In Vitro Model of a Cortical Microcircuit, Cerebral Cortex, Oxford University Press

Gupta Pratiksha, Mehra Rajesh, 2016, Low Power Design of SR Flip Flop Using 45nm Technology, IOSR Journal of VLSI and Signal Processing (IOSR-JVSP) Volume 6, Issue 2, Ver. I (Mar. -Apr. 2016), PP 54–57

Harris Kenneth D. et. al. 2001, Temporal Interaction between Single Spikes and Complex Spike Bursts in Hippocampal Pyramidal Cells, Neuron, Vol. 32, 141–149, October 11, 2001 Hippocampome.org, RRID:SCR_009023

Izaki Yoshinori, Takita Masatoshi and Tatsuo Akema, 2008, Specific role of the posterior dorsal hippocampus–prefrontal cortex in short-term working memory, European Journal of Neuroscience, Vol. 27, pp. 3029–3034, 2008

Jochems, Arthur and Yoshidam, Motoharu, Persistent firing supported by an intrinsic cellular mechanism in hippocampal CA3 pyramidal cells, European Journal of Neuroscience, Vol. 38, pp. 2250–2259, 2013

Kaminski Jan, Sullivan Shannon, Chung Jeffrey M et al., 2017, Persistently active neurons in human medial frontal and medial temporal lobe support working memory, Nature Neuroscience

Komendantov Alexander O., Siva Venkadesh, et. al., 2019, Quantitative firing pattern phenotyping of hippocampal neuron types, Scientific Reports, 17915 (2019)

Le Duigou Caroline, XEtienne Savary, Dimitri M. Kullmann and Richard Miles, 2015, Induction of Anti-Hebbian LTP in CA1 Stratum Oriens Interneurons: Interactions between Group I Metabotropic Glutamate Receptors and M1 Muscarinic Receptors, The Journal of Neuroscience, October 7, 2015 • 35(40):13542–13554

Masse Nicolas Y., Guangyu R. Yang, H. Francis Song, Xiao-Jing Wang and David J. Freedman, 2019, Circuit mechanisms for the maintenance and manipulation of information in working memory, Nature Neuroscience volume 22, pages 1159–1167 (2019)

Mayford Mark, Steven A. Siegelbaum and Eric R. Kandel, 2012, Synapses and Memory Storage, Cold Spring Harb Perspect Biol 2012;4:a005751

McCormick David A., Shu Yousheng, Hasenstaub Andrea, et al., 2003, Persistent Cortical Activity: Mechanisms of Generation and Effects on Neuronal Excitability, Cerebral Cortex, Oxford University Press

Papoutsi Athanasia, Sidiropoulou Kyriaki, et al. 2013, Induction and modulation of persistent activity in a layer V PFC microcircuit model, Frontiers in Neural Circuits

Paré D, Collins DR., 2000, Neuronal correlates of fear in the lateral amygdala: multiple extracellular recordings in conscious cats. J Neurosci. 2000 Apr 1;20(7):2701–10. doi: 10.1523/JNEUROSCI.20-07-02701.2000. PMID: 10729351; PMCID: PMC6772231.

Pinsky Paul F. and John Rinzel, 1994, Intrinsic and Network Rhytmogenesis in a Reduced Traub Model for CA3 Neurons, Journal of Computational Neuroscience, 1, 39–60 (1994),

Rahman, Jamilur, and Berger, Thomas, Persistent activity in layer 5 pyramidal neurons following cholinergic activation of mouse primary cortices, European Journal of Neuroscience, Vol. 34, pp. 22–30, 2011

Rosenberg Nadia, Urs Gerber and Jeanne Ster, 2016, Activation of Group II Metabotropic Glutamate Receptors Promotes LTP Induction at Schaffer Collateral-CA1 Pyramidal Cell Synapses by Priming NMDA Receptors, The Journal of Neuroscience, November 9, 2016 • 36(45):11521–11531 • 11521

Sirota Anton et. al., 2008, Entrainment of neocortical neurons and gamma oscillations by the hippocampal theta rhythm, Neuron. 2008 November 26; 60(4): 683–697. doi:10.1016/j.neuron.2008.09.014.

Wang Hui, Alvaro O. Ardiles et. al., 2016, Metabotropic Glutamate Receptors Induce a Form of LTP Controlled by Translation and Arc Signaling in the Hippocampus, The Journal of Neuroscience, February 3, 2016 • 36(5):1723–1729 • 1723

Wang XZemin, Ryan Neely and XCarole E. Landisman, 2015, Activation of Group I and Group II Metabotropic Glutamate Receptors Causes LTD and LTP of Electrical Synapses in the Rat Thalamic Reticular Nucleus, The Journal of Neuroscience, May 13, 2015 35(19):7616–7625

Xu J, Clancy CE (2008) Ionic Mechanisms of Endogenous Bursting in CA3 Hippocampal Pyramidal Neurons: A Model Study. PLOS ONE 3(4): e2056. https://doi.org/10.1371/journal.pone.0002056

